# Low Rate Hippocampal Delay Period Activity Encodes Behavioral Experience

**DOI:** 10.1101/2023.01.09.523199

**Authors:** Markos Athanasiadis, Stefano Masserini, Li Yuan, Dustin Fetterhoff, Jill K Leutgeb, Stefan Leutgeb, Christian Leibold

## Abstract

Remembering what just happened is a crucial prerequisite to form long-term memories but also for establishing and maintaining working memory. So far there is no general agreement about cortical mechanisms that support short-term memory. Using a classifier-based decoding approach, we report that hippocampal activity during few sparsely distributed brief time intervals contains information about the previous sensory motor experience of rodents. These intervals are characterized by only a small increase of firing rate of only a few neurons. These low-rate predictive patterns are present in both working memory and non-working memory tasks, in two rodent species, rats and Mongolian gerbils, are strongly reduced for rats with medial entorhinal cortex lesions, and depend on the familiarity of the sensory-motor context.

## Introduction

The behavioral relevance of a recently experienced event is not necessarily apparent during or shortly after it occurs. Nevertheless, it has to be maintained in memory for some time to potentially associate it with a subsequent reward or punishment – a requirement known as *temporal credit assignment problem* (Sutton, 1984; Sutton and Barto, 2018). Reinforcement learning (RL) solves this problem by both integrating a reward prediction value over time (Schultz et al., 1997) and propagating it through state space via an eligibility trace (Sutton and Barto, 1981), which might be implemented on the synaptic level (Päpper et al., 2011; He et al., 2015). While the neural mechanisms underlying reward prediction in the ventral tegmental area are exceptionally well investigated, the cortical activity that provides the experience-specific drive for tegmental RL processes largely remained unresolved. Potential cortical mechanisms to maintain short-term memory are persistent cortical activity (Egorov et al., 2002) and short-term synaptic plasticity (Mongillo et al., 2008; Leibold et al., 2008). A further potential mechanism are hippocampal time cells (MacDonald et al., 2011), however, they require the animal to be engaged in an active working memory task (Pastalkova et al., 2008) and are not necessarily content specific (Sabariego et al., 2019). Moreover, animals with bilateral lesions of the medial entorhinal cortex (mEC) show a behavioral deficit in spatial working memory but time cell activity in the delay period did not seem to be impaired (Sabariego et al., 2019). Since short-term memory is a prerequisite for working memory, we reckoned that mEC lesions might already affect the former and searched for potential impairments of the delay activity in the animals with mEC lesions, which may not be reflected in time cell activity. We, indeed, were able to identify activity correlates of behavioral performance differences between control rats and mEC-lesioned rats (Sabariego et al., 2019): activity from CA3 in control animals was more predictive of the previous behavioral trial than activity from CA3 in mEC-lesioned animals. The informative components of the activity were carried by only few cells that fired few additional spikes. In addition to the rat data, we also assessed CA1 activity of Mongolian gerbils (Fetterhoff et al., 2021) during a reward consumption period that did not require to maintain working memory and found identical results than for control rats, suggesting that the informative low-rate activity patterns are not working-memory dependent, but may constitute a hippocampal trace of cortical short-term memory processing.

## Results

To explore the information content of hippocampal activity during waiting periods, we examined two data sets in which animals performed different behavioral tasks. In a first data set, two groups of rats (with and without bilateral mEC lesions) were trained on a spatial alternation task with a variable waiting period between trials, in which they needed to maintain a working memory of their previous behavioral choice (Supplementary Figure S1A). Here, we only focused on sessions with 60 s long delay periods. Previously, it was shown that in animals with mEC lesions task performance is degraded (Sabariego et al., 2019) but it remained unclear whether this behavioral finding is reflected in hippocampal activity during the delay period. In a second data set Mongolian gerbils were trained to run on two mazes in virtual reality (distinguished by left and rightward turns and a turn-direction specific set of visual cues; Supplementary Figure S1B), that were selected in random order (such that no information about the future can be represented in inter trial intervals and animals had no requirement of working memory), and had a 20 second pause between trials during which animals received a reward (Fetterhoff et al., 2021). We compared two types of virtual mazes, a familiar configuration of visual cues in which the animals have been trained on the task, and a “swapped” maze in which visual cues are presented in association with the other turn direction introducing sensory conflicts, while the animals kept performing the same task.

### Decoding Performance

In the original analysis for the delay activity in the rat data sets (Sabariego et al., 2019), a linear (support vector) classifier was unable to distinguish whether preceding trials had left and right turns when population vectors were constructed with *L* = 1 s binning. Here, we repeated the analysis with a shorter time interval *L* = 100ms matching the typical duration of population bursts and a linear neural network to predict the left/right label of the trial preceding the delay period in all six groups of experiments. Correct classification rates (CCR) were slightly but significantly above chance (see example in Figure 1A) in a fraction of rat recording sessions that exceeded randomness (according to binomal tests, see Figure caption) except for CA1 recordings from rats with mEC lesions (Figure 1B). Original virtual reality mazes could also significantly be decoded from delay activity (Figure 1B, LR maze), but swapped mazes with sensory conflicts could not (Figure 1B, L*R* maze). Since we observe predictions of previous trial labels, even in the gerbil data set without a working memory task, we reason that the activity does not specifically reflect working memory. Nevertheless the activity may underlie the establishment of working-memory although differences between mEC-lesioned and control rats do not yet reach significance at this level of analysis.

**Figure 1:**
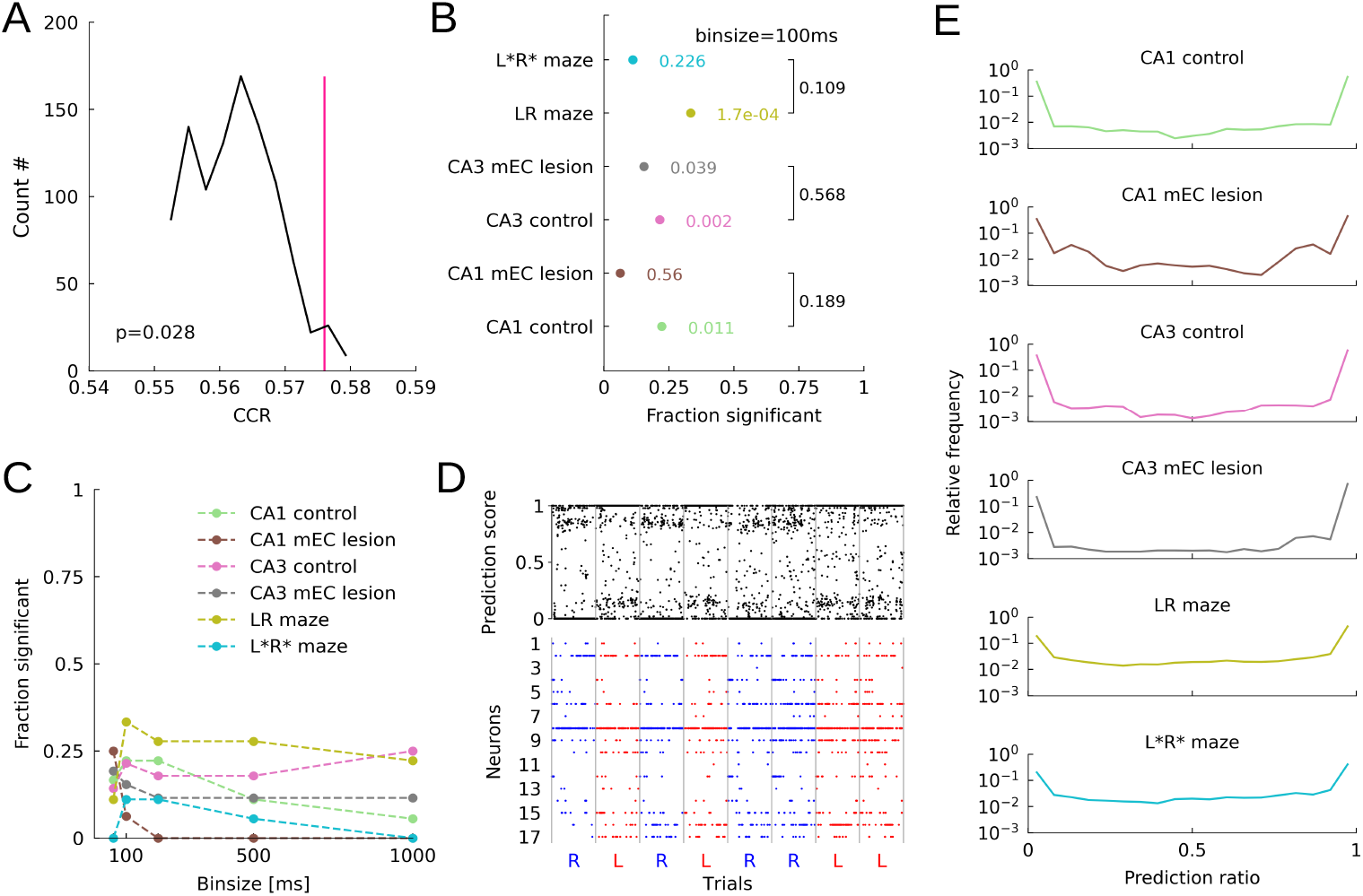
Decoding performance of linear ANN. (A) Correct classification rate (CCR; pink) for an example CA1 recording from a control rat (rat 3903/day 1/session 2) and distribution of CCRs for label shuffles (black). Despite being small, the CCR is significant (p value as indicated; 1000 shuffles; 100 cross validation iterations). (B) Fraction of sessions for which the permutation test from A was significant for each of the six groups of experiments (p values from binomial tests; CA1 control: 4/18; CA3 control: 6/28; RL maze: 6/18). mEC lesions in rats and unfamiliar arrangement of visual landmarks in gerbil data lead to decreased decoding (p values as indicated; CA1 mEC lesion: 1/16; CA3 mEC lesion: 4/26; L*R* maze: 2/18 Chi squared test for homogeneity, rat CA1: *χ*^2^ = 1.723, *n*_1_ = 4, *n*_2_ = 1; rat CA3: *χ*^2^ = 0.326, *n*_1_ = 6, *n*_2_ = 4; gerbil CA1: *χ*^2^ = 2.571, *n*_1_ = 6, *n*_2_ = 2; See Table 1). (C) The fraction of significantly decodable sessions decreases for larger time intervals *L* in CA1 data sets but remains at a constant level in CA3 data sets. (D) Prediction score (PS) for an example session from CA1 of a rat with mEC lesion (Rat 3928/day 2/session 1) using *L* = 100*ms* time intervals (top) and spike raster plot (bottom) from a 60 s delay period succeeding left- and right-ward trials (red and blue, respectively). (E) Distributions of the PS for all groups of experiments only exhibit a small bias towards 1.

The fraction of significant sessions was generally highest for *L* = 100 ms intervals (except in CA1 recordings from MEC-lesioned animals, where the fraction of significant sessions was maximal for 50 ms binning) and decreased with larger bin sizes (Figure 1C; except for CA3 in control rats) indicating that, at least in CA1, the information about the previous turning direction is mostly carried by short-term correlations The finding that CA3 activity even for *L* = 1 s is significantly predictive contradicts previous reports in (Sabariego et al., 2019) and may arise due to a linear neural network classifier instead of a linear support vector machine and/or different preprocessing (scaling). Further insight into the predictive activity patterns (see Section “Low Rate Relevant Time Bins”), will further explain differences between CA1 and CA3 results.

**Table 1:** Figure 1B,Chi squared test for homogeneity, Test statistics, p values, degrees of freedom (df)

| Condition | Test statistics | p values | degrees of freedom(df) |
| --- | --- | --- | --- |
| CA1 control<br>CA1 mEC lesion | 1.723 | 0.189 | $n_1 = 4, n_2 = 1$ |
| CA3 control<br>CA3 mEC lesion | 0.326 | 0.568 | $n_1 = 6, n_2 = 4$ |
| CA1 LR maze<br>CA1 L*R* maze | 2.571 | 0.109 | $n_1 = 6, n_2 = 2$ |

To better understand what activity features the classifiers use to distinguish past experiences, we first computed a prediction score (PS) for every time bin. The PS measures the fraction of repetitions in which a population vector from a particular time bin yielded a correct prediction during testing (see Methods). One representative example session (Figure 1D) reflects a general directional bias (here “left”) of the classifier, i.e., the prediction outcomes tend to favour the label “left” independent of the real trial label if no information seems available in the spiking pattern. The above chance performance of the classifier on average (Figure 1E) is reflected in PS distributions with a peak at 1 only slightly exceeding the peak at 0.

One possible explanation for the low average CCR values and the small bias in PS that is in accordance with the general increase in prediction for lower bin sizes *L* is to assume that the informative neural signatures occur only in few brief time intervals. A natural guess would therefore be to investigate the association between intervals of high PS and awake sharp-wave ripple (SWR) events, since they are of about 100 ms length, are generally thought to support planning (Jadhav et al., 2012; Shin et al., 2019), and the incidence rates are affected by functional mEC inputs (Chenani et al., 2019).

We tested this conjecture for the control data sets from rats (CA1 and CA3) performing a spatial working memory task, by correlating the local field potential (LFP) power in different frequency bands with the prediction scores of the classifier in 100ms bins. We, however, did not find any consistent correlation between spectral bands and prediction (Figure 2A) by standard multilinear regression. Only 1 of 4 significant CA1 session and 1 of 5 significant CA3 session showed overall significant linear relation (ANOVA) with 3 of 9 (=4 CA1+ 5 CA3) individual tests showing significance in the theta band and 2 of 9 in the ripple band. Wilcoxon signed-rank tests for non-zero regression weights across sessions (black circles in Figure 2B) did show no significant results, suggesting that overall dependencies of prediction scores on LFP must be weak. This conclusion was further corroborated by inconsistent significance of correlations between LFP power and prediction scores when data were pooled over all sessions. Pooled CA1 prediction scores, were significantly modulated with ripple power, but not the pooled CA3 prediction scores (red circles in Figure 2B). We visualized the best candidate correlations (CA1 theta and ripple) as scatter plots, which revealed that the significant linear regression of the pooled prediction scores may only explain a negligibly small part of the variance (Figure 2C). With this observed lack of clear correlation we rule out that successful decoding mostly relies on SWR or any other LFP-related activity pattern.

**Figure 2:**
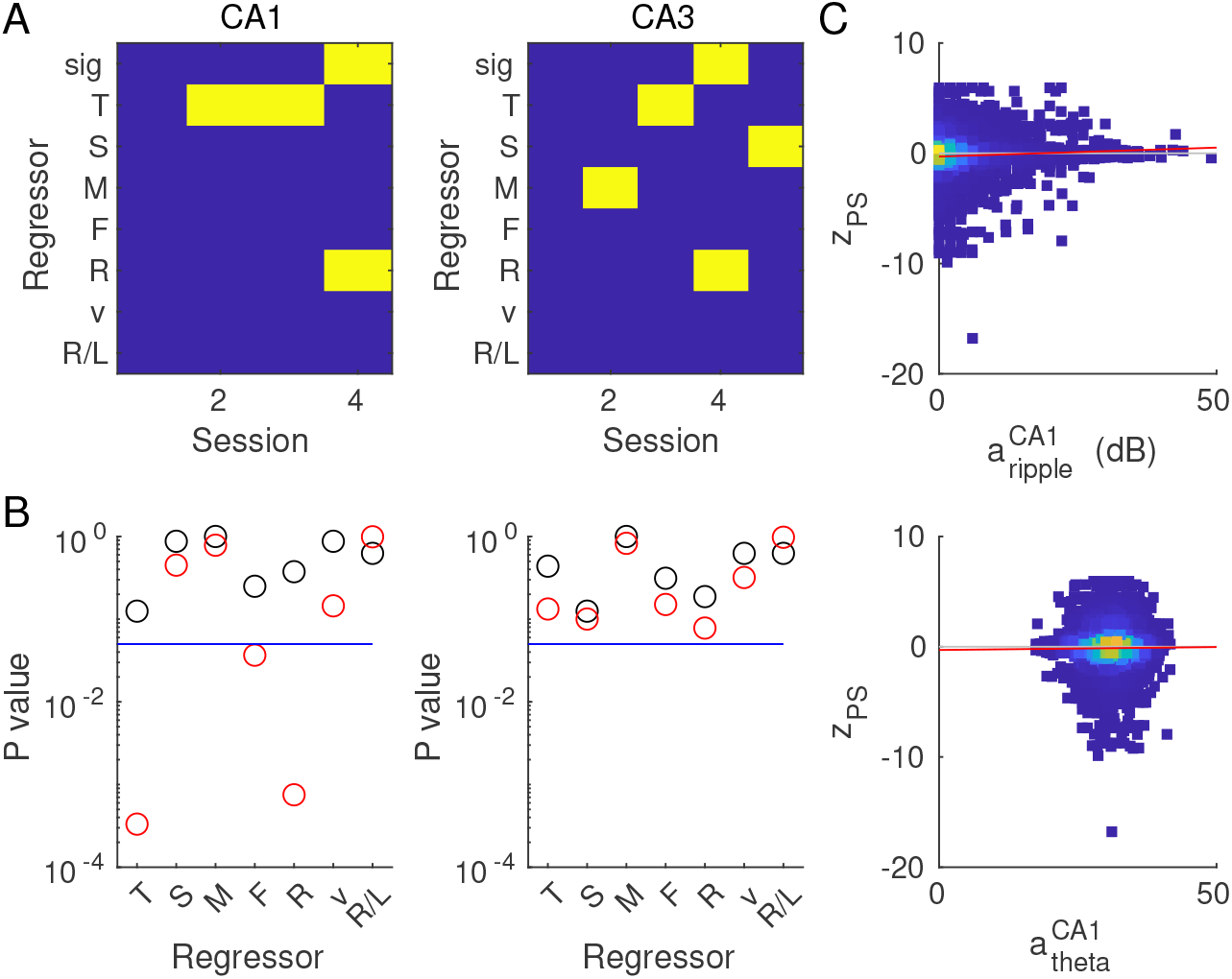
Lack of clear correlation between z-scored prediction score (PS) and LFP power (in dB) for the 9 significant control sessions. (A) Significance (p-value < 0.05: yellow; p-value > 0.05: blue) of general linear model fits of the z-scored PS using the regressors theta power (T, 6–11 Hz), slow- (S, 30–50 Hz), mid- (M, 55–90 Hz), fast-(F, 95–140 Hz) gamma, and ripple (R, 150–250 Hz) power, speed (v), and label (R/L) for CA1 (Left) and CA3 (right) recordings for control rats. Significance for the whole model fit (sig) is obtained from F-statistics, significance for the *β* values from T-statistics. Numerical values for test statistics and p values are provided in the Supplementary Table 1. (B) P values for fits to pooled data (T-statistics, red) and Wilcoxon tests on regression coefficients (*β*-values) of the individual sessions being different from zero (black). (C) Scatter (density) plots for z-scored PS vs. regressors theta power (bottom) and ripple power (top) with regression line (red).

### Most Informative Directions

To directly identify the neuronal basis of the prediction scores of the classifier we intended to visualize its decision boundary, i.e., to identify the neural ensembles that are specific to the previous experience of the animal. To do so, we applied adversarial attack techniques from machine learning (see Methods) (Rauber et al., 2017; Goodfellow et al., 2014) that move the population vector constructed from a specific time point to a position close to the classification boundary (Figure 3A,B). From this set of boundary positions we then constructed most informative directions (MIDs, orange vector in Figure 3B) as clusters of orthogonal vectors to the boundary. The method outperforms estimating the weight vector by bootstrapping the training process on multiple subsamplings for low signal strength (Supplementary Figure S2A-C).

**Figure 3:**
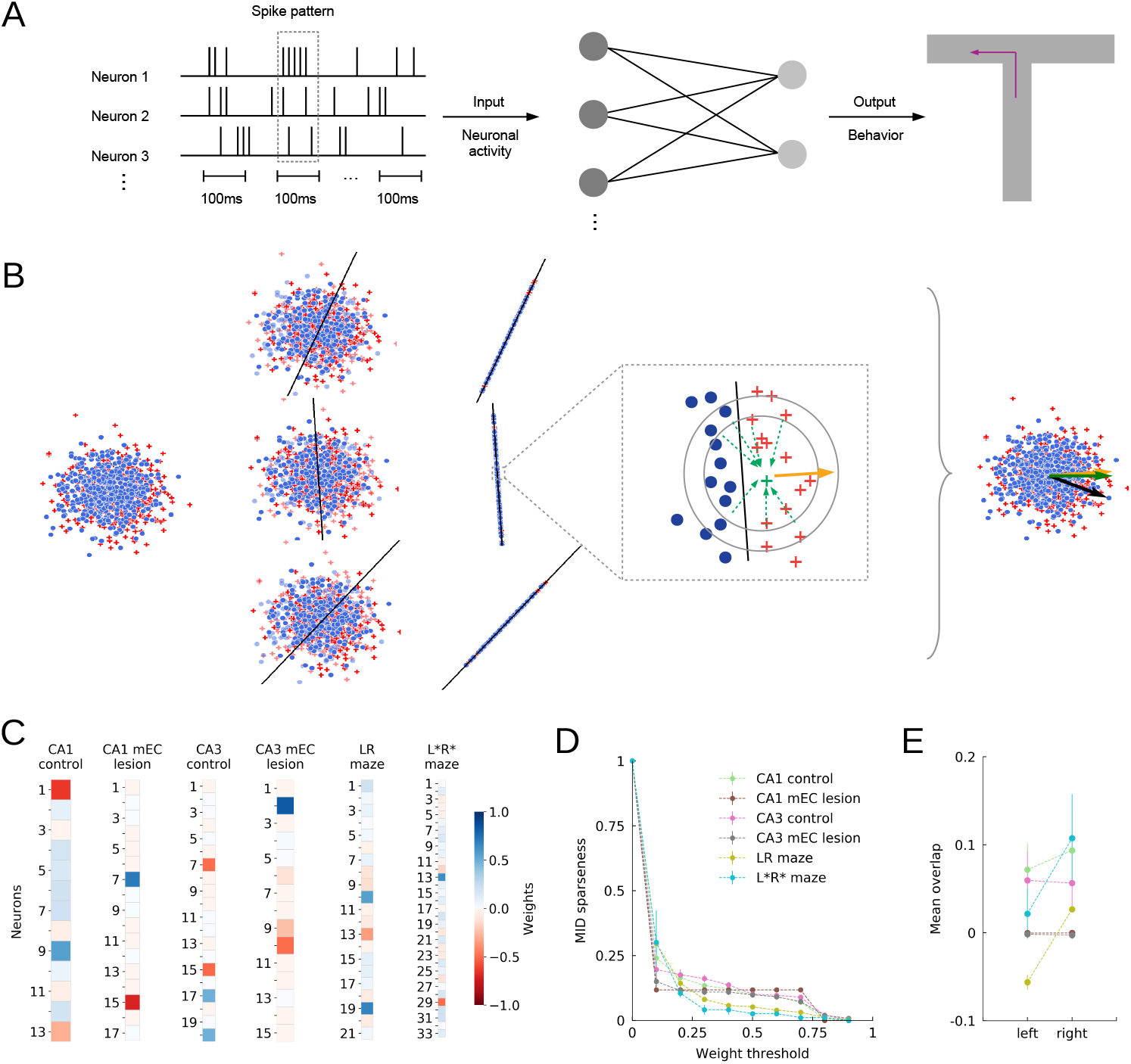
Most informative directions (MIDs). (A) Schematic of the decoding process using a linear neural network and population vectors with binary labels from behavior. (B) Illustration of the MID identification process (Methods). A linear classifier is trained using a 2-fold cross validation scheme. Adversarial attack methods are employed to move data points close to the decision boundary. MIDs are then identified by clustering (DBSCAN) locally orthogonal vectors (gold). Results are grouped, sorted and evaluated over all cross validation iterations (100 bootstraps). (C) MIDs from example sessions for all groups of experiments (CA1 control rat 3906/day 1/session 2;CA1 mEC-lesioned rat 3928/day 2/session 1; CA3 control rat 3958/day 3/session 2; CA3 mEC-lesioned rat 3903/day 1/session 2;RL maze gerbil 2783/day 1; R*L* maze gerbil 2784/day 3). Saturated colors indicate neurons which contribute more strongly to the decision boundary. (D) Fraction of MID weights (sparseness) exceeding a certain threshold. (E) Mean overlaps of population vectors with MIDs for “left” and “right” labelled trials.

Examples for MIDs from all 6 data sets are shown in Figure 3C, indicating only few active neurons (saturated colors) to contribute to the decision of the classifier. Varying the weight threshold to obtain heuristic sparseness estimate reveals that only about 20% of the neurons (that were active in the delay period) may contribute to the classification performance (Figure 3D).

To test whether the obtained MIDs indeed identify functionally relevant dimensions, we assigned overlap values 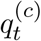 with all the MIDs (identified by *c*) to the population vectors (identified by time index *t*). If the sign of *q* correlates with the decision performance, we would consider the MID informative. However, we only find such a sign change to occur in the control gerbil data set (Figure 3E), suggesting that averaging over all time bins probably blurs the signal and thus proceeded with restricting our analysis to only those “relevant” time bins which we suspect to be most informative.

### Low Rate Relevant Time Bins

To identify relevant time bins, we compared the overlap values *q* with the shuffle distribution (see Methods; Supplementary Figure S3) to find above-chance overlap with the MID. Time bins for which *q*_*t*_ was below the 2.5 percentile of the shuffle distribution (significantly negative overlap) were considered to be predictive for “left” labels, time bins for which *q*_*t*_ was above the 97.5 percentile (significantly positive overlap) of the shuffle distribution were considered to be predictive for “right” labels. Figure 4A depicts two examples of spike patterns from the relevant time bins. These examples are typical (see further examples in Supplementary Figure S4), in that the firing rate in relevant bins of mostly only one neuron considerably exceeds its firing rate in the non-relevant bins, and this neuron gets the largest load of the MID in positive (right) and negative (left) direction. We also observe general modulations of firing rates across trials with some trials having increased activity in all neurons.

**Figure 4:**
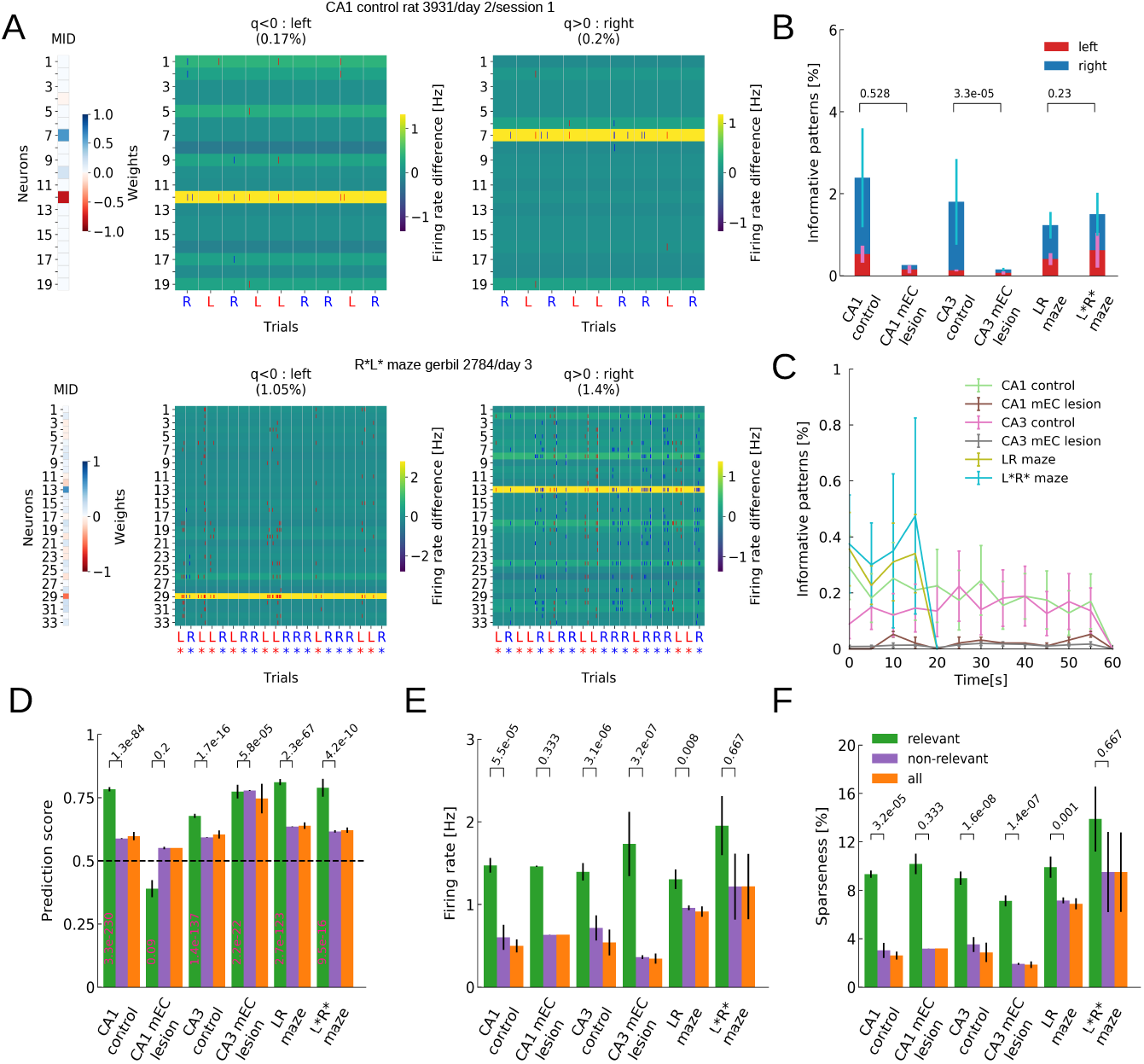
Relevant time bins. (A) MIDs (left) and spike raster plots (right) only for relevant time bins of two (rows) example sessions (top: rat 3931/day 2/session 1; bottom: gerbil 2784/day 3). Spikes are colored (red/blue) according to trial labels. The background colors indicate the differences in firing rate between relevant and non-relevant bins of the individual neurons. Percentages on top reflect fractions of bins identified as relevant. (B) Mean percentage of relevant time bins for each condition with respect to “left”- (red) and “right”-ward (blue) turns. mEC-lesioned rats showed significantly fewer relevant time bins (Mann-Whitney U test; rat CA1: *U* = 23.5, rat CA3: *U* = 487.5, gerbil CA1: *U* = 8.5; See Table 5). (C) Distribution of relevant time bins across the delay periods for all sets of experiments. (D) The prediction scores for relevant time bins are significantly above chance for all sets of experiments except for the CA1 in mEC-lesioned animals (Wilcoxon test; See Table 6). Prediction scores of relevant time bins are significantly larger than the non-relevant ones in all sets of experiments apart from the mEC-lesioned animals (Mann-Whitney U test; see Table 7) (E) Significantly higher firing rates are observed in relevant vs. non-relevant time bins for all sets of experiments apart from the CA1 of lesioned rats and the unfamiliar L*R* mazes in gerbil CA1 (Mann-Whitney U test, see Table 8). (F) Significantly higher fraction of active cells are observed in relevant vs. non-relevant time bins for all sets of experiments apart from the CA1 from lesioned rats and the unfamiliar L*R* mazes in gerbils (Mann-Whitney U test, see Table 9).

These two examples are also typical, in that only a small fraction of time bins turned out to be relevant in general (Figure 4B) with most relevant bins (2.4%) in CA1 data from control rats. Rats with mEC lesions showed particularly low fractions of relevant bins with the difference between control and lesioned animals reaching significance only for CA3 recordings (Mann-Whitney U rank test). This finding suggests the mEC supports the expression of brief periods of informative delay period activity that, at least in CA3, may reflect working memory performance. The sparsely interspersed relevant time bins thereby occur at similar rates across the delay period in all analysis groups (Figure 4C).

**Table 2:** Figure 2A, CA1, Test statistics, p values, degrees of freedom (df)

| rat/day/sess. | 3906/1/1 | 3906/1/2 | 3931/2/1 | 3931/3/2 |
| --- | --- | --- | --- | --- |
| ANOVA (F) | 1.08, 0.38 | 1.75, 0.093 | 1.93, 0.061 | 7.16, 1.6e-8 |
| T | 0.48, 0.64 | 2.05, 0.041 | 2.1, 0.032 | 1.50, 0.13 |
| S | -1.11, 0.27 | -1.38, 0.17 | 1.35, 0.18 | -0.50, 0.62 |
| M | -0.15, 0.89 | 0.34, 0.74 | 0.68, 0.50 | -0.90, 0.37 |
| F | -1.13, 0.26 | -1.42, 0.16 | 0.44, 0.66 | -1.78, 0.075 |
| R | 1.91, 0.056 | 1.18, 0.24 | -1.36, 0.17 | 6.53, 7.7e-11 |
| v | 0.98, 0.33 | 0.88, 0.39 | 1.68, 0.093 | -1.61, 0.11 |
| R/L | -0.10, 0.92 | 0.19, 0.86 | -0.08, 0.94 | 0.24, 0.82 |
| df | 4774 | 5381 | 4779 | 3584 |

**Table 3:**
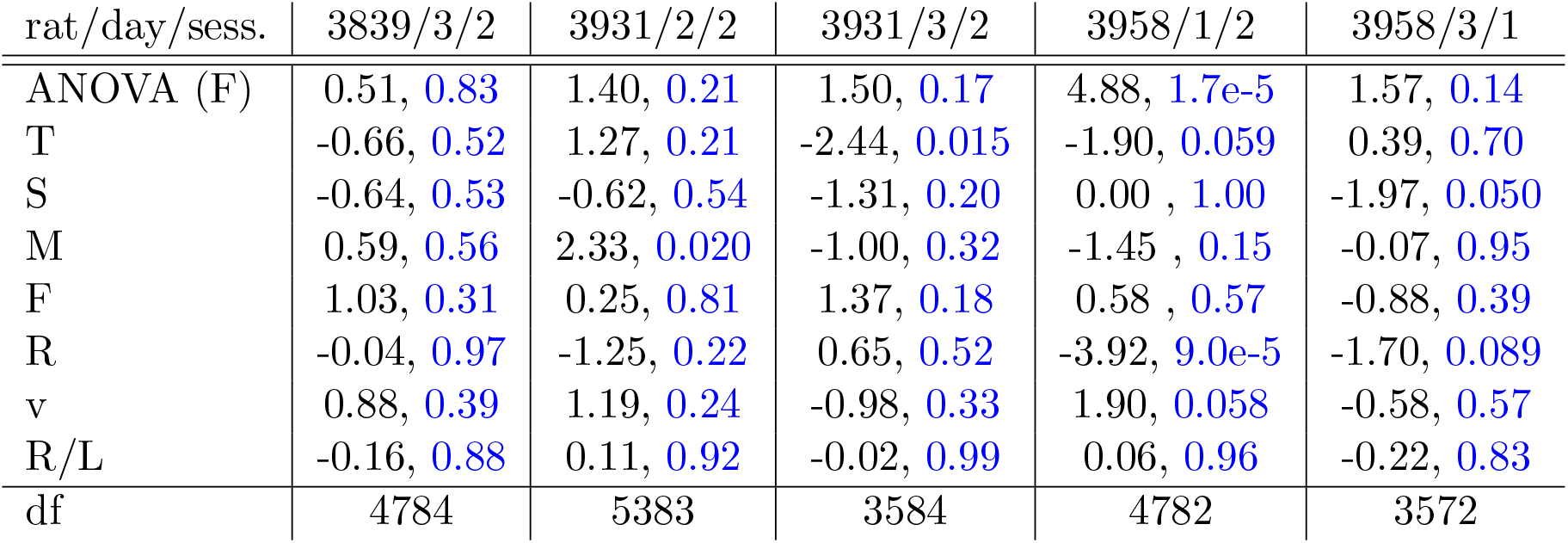
Figure 2A, CA3, Test statistics, p values, degrees of freedom (df)

**Table 4:**
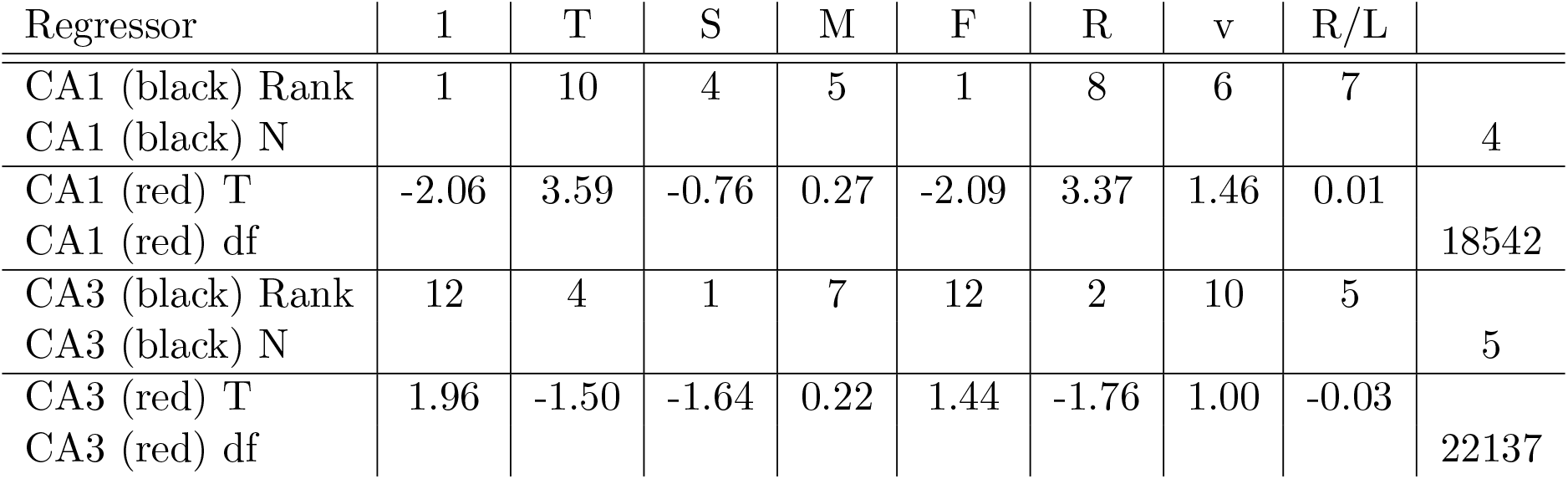
Figure 2B, Test statistics, (N,df)

**Table 5:**
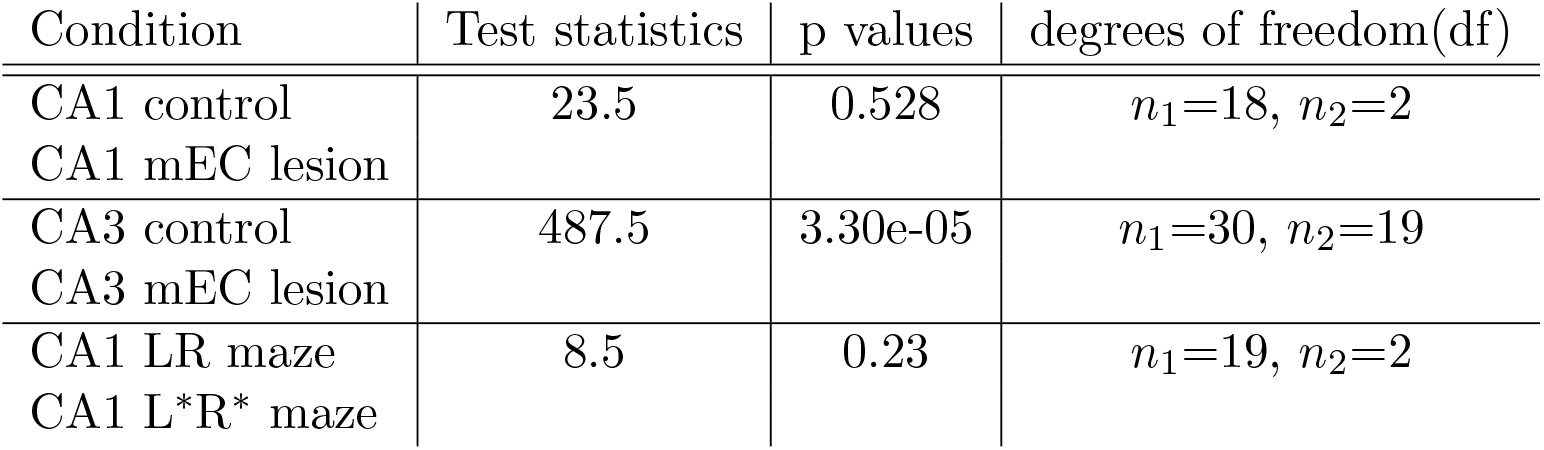
Figure 4B,Mann-Whitney U test, Test statistics, p values, degrees of freedom (df)

| Condition | Test statistics | p values | degrees of freedom(df) |
| --- | --- | --- | --- |
| CA1 control | 23.5 | 0.528 | $n_1=18, n_2=2$ |
| CA1 mEC lesion |  |  |  |
| CA3 control | 487.5 | 3.30e-05 | $n_1=30, n_2=19$ |
| CA3 mEC lesion |  |  |  |
| CA1 LR maze | 8.5 | 0.23 | $n_1=19, n_2=2$ |
| CA1 L*R* maze |  |  |  |

**Table 6:** Figure 4C, Wilcoxon test, Test statistics, p values, degrees of freedom (df)

| Condition | Test statistics | p values | degrees of freedom(df) |
| --- | --- | --- | --- |
| CA1 control | 32.394 | 3.28e-230 | 2127 |
| CA1 mEC lesion | -1.697 | 8.95e-4 | 25 |
| CA3 control | 24.968 | 1.37e-137 | 3215 |
| CA3 mEC lesion | 9.7334 | 2.164e-22 | 161 |
| CA1 LR maze | 23.615 | 2.714e-123 | 939 |
| CA1 L*R* maze | 8.033 | 9.50e-16 | 120 |

**Table 7:** Figure 4C, Mann-Whitney U test, Test statistics, p values, degrees of freedom (df)

| Condition | Test statistics | p values | degrees of freedom(df) |
| --- | --- | --- | --- |
| CA1 control | 11.276e7 | 1.316e-84 | $n_1=2127, n_2=86685$ |
| CA1 mEC lesion | 10.304e4 | 0.2 | $n_1=25, n_2=9575$ |
| CA3 control | 26.615e7 | 1.705e-16 | $n_1=3215, n_2=153987$ |
| CA3 mEC lesion | 72e5 | 5.762e-05 | $n_1=161, n_2=103639$ |
| CA1 LR maze | 46.367e6 | 2.340e-67 | $n_1=939, n_2=74943$ |
| CA1 L*R* maze | 62.289e4 | 4.24e-10 | $n_1=120, n_2=7837$ |

**Table 8:** Figure 4D, Mann-Whitney U test, Test statistics, p values, degrees of freedom (df)

| Condition | Test statistics | p values | degrees of freedom(df) |
| --- | --- | --- | --- |
| CA1 control | 290.0 | 5.473e-05 | 18 |
| CA1 mEC lesion | 4.0 | 0.333 | 2 |
| CA3 control | 766.0 | 3.092e-06 | 30 |
| CA3 mEC lesion | 356.0 | 3.213e-07 | 19 |
| CA1 LR maze | 272.0 | 0.008 | 19 |
| CA1 L*R* maze | 3.0 | 0.667 | 2 |

**Table 9:** Figure 4F, Mann-Whitney U test, Test statistics, p values, degrees of freedom (df)

| Condition | Test statistics | p values | degrees of freedom(df) |
| --- | --- | --- | --- |
| CA1 control | 294.0 | 3.168e-05 | 18 |
| CA1 mEC lesion | 4.0 | 0.333 | 2 |
| CA3 control | 833.0 | 1.554e-08 | 30 |
| CA3 mEC lesion | 361.0 | 1.443e-07 | 19 |
| CA1 LR maze | 291.0 | 0.001 | 19 |
| CA1 L*R* maze | 3.0 | 0.667 | 2 |

Despite PS over all time bins only had a tiny bias towards 1, prediction scores in the relevant bins are very clearly and significantly above chance (Wilcoxon test; see Table 6) for all data sets except CA1 recordings from lesioned rats. Also PS in relevant bins were significantly larger (Mann-Whitney U rank test; see Table 7) than in non-relevant bins except for the data sets from mEC-lesioned animals (Figure 4D). These findings indicate that MIDs provide a handle for identifying predictive neural activity except in the two data sets from lesioned animals, possibly because there are just too little relevant time bins (Figure 4B).

The number of spikes contributing to the above chance prediction is very small as indicated by firing rate (Figure 4E) and sparseness (Figure 4F), but most data sets exhibit significantly increased firing rate and fractions of active cells in relevant bins as compared to non-relevant bins (Mann-Whitney U rank test; see Tables 8, 9), indicating that indeed few brief intervals of slightly enhanced activity carry the behavioral information. In this context, we also revisited the unexpectedly large predictability of CA3 activity in long time bins of *L* = 1 s (Figure S5), and found a lower firing rate in the relevant bins and a relatively lower difference (as compared to CA1) in sparseness between relevant and non-relevant time bins. This indicates that predictive activity in CA3 seem to be dispersed over longer time periods than in CA1.

## Discussion

We examined the short-term memory content of hippocampal CA1 and CA3 activity for rats during a delayed spatial alternation task and CA1 activity for Mongolian gerbils, after navigating virtual reality mazes, during reward consumption in inter-trial intervals. The recorded activity of past experience was decoded applying a linear neural network to population vectors from time bins of length of 100 ms. To directly identify the neuronal basis of the prediction accuracies we visualized the decision boundary using adversarial attacks, and subsequently identified most informative neuronal ensembles in terms of vector clusters orthogonal to the decision boundary. Applying these neuronal ensembles to the recorded activity we were able to extract the activity patters most related to previous behavior. Few neurons (about 20% of those that were active in the delay periods) and few time bins (2%) seem to be contributing to the classification task with relatively low firing rates (about 2.5 Hz) across all experimental conditions. We reasoned that this may indicate that the recent past is encoded with sparsely dispersed spikes. Since the amount of informative activity patterns was reduced at least in CA3 of lesioned rats, our results suggest that this low-rate activity may support working memory processes.

The medial entorhinal cortex (mEC) provides the hippocampus with spatial information during foraging and navigation tasks (Hafting et al., 2005; Gil et al., 2018). Previous analysis observed that mEC is necessary for control level working memory performance (Sabariego et al., 2019), despite only limited effects on hippocampal place fields (Hales et al., 2014; Schlesiger et al., 2015) and sequence replay (Chenani et al., 2019). Our results suggest an additional mEC-dependent mode of activity that appears to hold information related to previous experience, which entails a relatively low number of active cells and spikes. A further decline of predictivity was observed in gerbils navigating through virtual environments with conflicting sensory-motor context, suggesting that particularly sensory information via the mEC may be a main driver for the informative low rate activity patterns.

Because the observed brief periods of informative activity are sparse and random in time, potential mechanisms that may give rise to them are unlikely to consist of local persistent neural firing generated by positive self-feedback (Fransén et al., 2006). Synfire chains (Abeles, 1991) that propagate through multiple brain areas, however, cannot be excluded, but would require that similar temporally correlated activity signatures would be visible in other limbic brain areas. Particularly the medial prefrontal cortex with its direct hippocampal innervation, however, may lack such informative activity (Böhm and Lee, 2020). An alternative mechanism to store short-term memory is synaptic short-term dynamics (Mongillo et al., 2008; Leibold et al., 2008). The synapse-specific depression and facilitation states may maintain specific behavioral information for time scales up to few seconds, however, these states would need to be refreshed every few seconds to bridge intervals of several tens of seconds as in the currently investigated behavioral tasks. The low rate activity patterns described in this paper may implement such a refreshing mechanism.

Classifier-based decoding is a robust method to link specific features of neuronal activity to cognitive function and behavior. The existence of several well-tested classifier implementations that are straight-forward to analyze within the theoretical framework of hypothesis testing and cross validation is particularly convenient (Bishop, 2006) and makes them good candidates for decoding typically low signal to noise neuronal activity (Haynes and Rees, 2006; Norman et al., 2006; Lemm et al., 2011). The downside of classifier-based decoders is that they usually come as a black box, meaning that the neuronal activity features which are the most influential regarding a specific behavioral outcome are not readily observable. However, knowing these features, is pivotal for correlating neuronal ensembles to behavioral states. Here, we employ explainable artificial intelligence methods (Karimi et al., 2019), known as adversarial attacks, in order to sample the decision boundary of classifiers in an attempt to overcome their black-box nature (Doran et al., 2017). In doing so, we identify the most informative neuronal ensembles in terms of consistently appearing clusters of normal vectors relative to the decision boundary.

Classification of high-dimensional (multi-neuron) data with low signal to noise ratio and limited numbers of trials is usually best done with linear models, since more complex non-linear classifiers are prone to overfitting, exhibiting test performances that are drastically inferior to training performances (Bishop, 2006). Although linear classifiers may thus turn out superior in many of the real-world applications from an empirical risk minimization perspective, the true underlying generative processes may nevertheless be non-linear. Our attack-based approach provides a handle to uncover at least parts of the underlying non-linearities with a linear network, by multiple subsamplings for each of which we estimate the normal vector. Clustering of normal vectors from the many subsamplings and applying consistency measures allows to detect multiple considerably distinct clusters of normal vectors, and thus allows to effectively describe some of the non-linear structure in the data.

How to maintain information over time intervals of tens of seconds to minutes, and how to achieve this at low energetic costs, are key open problems in understanding the cortical basis of working memory. Particularly the energy constraint will restrict the neuronal activity correlates to be sparse and low-rate, properties that make them hard to find. Further new analysis approaches will be needed to identify such neural signatures, particularly also in correlation with behavioral measures (Schneider et al., 2022).

## Methods

### Electrophysiological Data Sets

Both of data sets included in our analysis have been previously published (Sabariego et al., 2019; Fetterhoff et al., 2021). Detailed descriptions of the experimental methods can be found in the original papers.

In brief, 15 male Long Evans rats were trained on the spatial alternation task (Sabariego et al., 2019) and randomly assigned to one of two groups, a group with nearly complete NMDA lesions of the medial entorhinal cortex (n = 7) and a control group (n = 8). After about 9 weeks of recovery, both groups of animals were implanted with tetrodes that were lowered until the CA1 or CA3 region. The behavioral task was performed using an 8-shaped maze (Figure S1A). Rats were trained until the performance reached 90% correct trials on two of three consecutive days. After that 30 trials with 60s delay were performed daily for each rat, for 14 days.

From the virtual reality task (Fetterhoff et al., 2021), we obtained data from six male Mongolian gerbils (Meriones unguiculatus) with tetrodes implanted to dorsal hippocampal CA1. Gerbils were trained to the task of running on a 620 cm long linear track consisting of three linear hallways separated by two 45^*o*^ corners. The animals were initially introduced on the two original maze types: R and L, each containing two right or two left turns and, each containing different images (Figure S2B). After learning original image-turning direction combinations, images were swapped in the middle and the last hallways (L* and R* mazes). During recording sessions, 20 randomly-ordered original mazes were presented before 20 randomly ordered swapped mazes. The VR system is described in greater detail in (Thurley et al., 2014).

### Population Vectors

Spikes of all *N* neurons recorded during the delay phases of a session are time-binned in *t* = 1, …, *T* intervals with bin size *L*. The number *T* of time bins is also called the size of the data set. Neurons without any spike are excluded from the data set (Table 10) resulting in population vectors 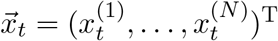. Each of the neurons is converted to the standardized space, by subtracting the mean neuron activity in a session and then dividing the difference by the standard deviation of the neuron activity,

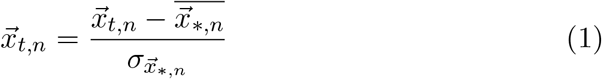

Each of the patterns 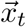 is assigned a label *l*_*t*_ = *±*1 according to the binary behavioral experience in the trial this pattern is obtained from. In our data sets, these binary labels distinguish rightward from leftward turns.

**Table 10:** Fraction of non-active cells (%)

|  | min – max | Mean |
| --- | --- | --- |
| CA1 control | 0.0 - 23.5 | 11.61 |
| CA1 mEC lesion | 5.0 – 5.0 | 5.00 |
| CA3 control | 0. - 66.7 | 18.1 |
| CA3 mEC lesion | 6.25 - 24.56 | 11.3 |
| CA1 LR maze | 0.0 | 0.0 |
| CA1 L*R* maze | 0.0 | 0.0 |

### Artificial data

#### Linear separation task

We generate a linearly separable data set of *t* = 1, …, *T* vectors

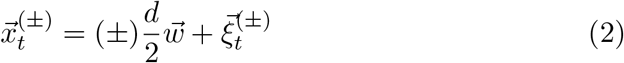

with labels *l*_*t*_ = ± 1. Here, *d* ≥ 0 denotes the signal along the ground truth direction 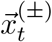 that is added (subtracted) to normal i.i.d random vectors 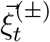. The dimension *n* of the vectors ranges between 2 and 100. The sparseness *s* = 1*/n* of the weight vector indicates only one active dimension, and the signal strength *d* varies between 0 and 10.

#### Artificial spiking model

To mimic random spiking activity we simulate *n* homogeneous Poisson processes with density *λ* (varying between 0.05 and 0.2) and construct *t* = 1, …, *T* population vectors from time bins of size 1 with balanced random labels *l*_*t*_ = *±*1.

A ground truth weight vector 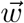 is then applied to a small subset *T* ^*′*^ = (1 − *p*_*fail*_) *T* of the available vectors from the positive subset (*l* = +1):

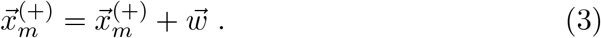

The sparseness *s* of the binary weight vector varies between 10 − 30%. The dimensions are varied between 10 and 50.

### Decoding

We train a binary classifier to distinguish the binary behavioral choices. Our specific choice of the classifier is a *linear* neural network, implemented in *PyTorch* (Paszke et al., 2019). The network consists of an input layer 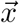, with one node for each of the active neurons we attempt to decode, and an output layer 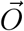 with two nodes, each dedicated to one of the binary labels. The output is computed using the softmax function

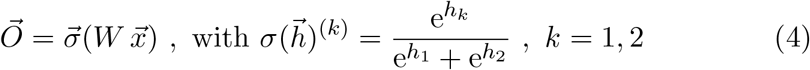

The training of the classifier minimizes the cross entropy loss function in the space of the 2 × *N* weight matrices *W* . Typically, supervised training occurs for 1000 consecutive epochs, with a learning rate 0.001 (see (Paszke et al., 2019)).

In order to ensure a less biased estimate of the model performance and to avoid overfitting, we employ a 2-fold crossvalidation process during which we generate 100 random separations into training and a testing subset of equal size *T/*2, in which the ratio between the two labels is kept as in the full data set. Applying the classifier on the test data in each of the 100 random separations yields a fraction of correct classifications. As correct classification rate (CCR) we define the mean of these 100 fractions.

We repeat the decoding for 1000 random shuffles of labels and thereby obtain 1000 CCRs (each averaged on 100 random separations into test and training sets) from which we construct the Null distribution that is used to assign a p value to the decoding performance as the percentile of the real CCR.

### Adversarial Attacks

To identify the separating hypersurface we ran two repetitions of the fast gradient sign method (FGSM) attack on each data point. The FGSM attack takes advantage of the gradient descent optimization of a neural network, and is executed via the Python-based package *Foolbox* (Rauber et al., 2017), which provides reference implementations of a variety of published state-of-the-art adversarial attacks (Goodfellow et al., 2014).

The attack maps each population vector 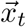 onto a different vector 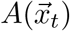 which is moved to the hemispace opposite to the separating hypersurface. We apply the attack process twice since then the resulting vectors 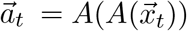 faithfully sample the classification boundary. After the two attacks a population vector 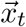 is thus associated with a vector 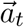 that is supposed to be proxy for the closest position on the separating hypersurface.

In order to avoid overfitting of the decision boundary, we repeat the computation of attack vectors 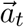 100 times using random subsamplings of the data set of size *T/*2 keeping the ratio of labels as in the full data set.

### Most Informative Directions

As most informative direction (MID) we define the direction in the space of population vectors 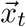 that is orthogonal to the decision boundary of the classifier. Since in general the decision boundary can be non-linear, MIDs are local and thus we expect that multiple MIDs can occur for any given dataset. It also needs to be noted that MIDs are a property of the data set and not the classifier. Thus even if we use a linear classifier to approximate parts of the (potentially non-linear) decision boundary, we may obtain multiple MIDs, depending on which part of the boundary is best matched by the current subsampling of the data set.

To obtain the orthogonal direction at one attack vector location 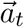, we compute a set of difference vectors 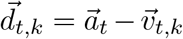 (Figure 3B, green arrows) with 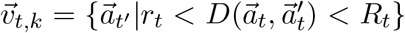 denoting subset of attack vectors in a ring-shaped vicinity of 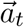. As a distance function *D* we use the Euclidean distance, with 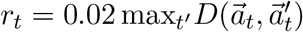 and 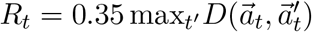.

The MIDs are then obtained by searching the directions 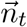 that minimize the squared scalar product to the distance vectors, i.e.,

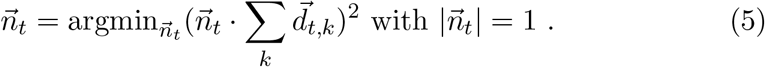

The minimization is equivalent to finding the Eigenvector 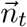 of the matrix 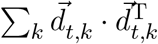 with the smallest Eigenvalue. Since the minimization problem in Eq. (5) is symmetric regarding multiplication with −1, we always choose 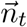 pointing into the +1 hemispace.

Finally, we apply the density-based spatial clustering algorithm (DB-SCAN from the Python package *scikit learn* (Pedregosa et al., 2011)), to all vectors 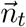 and to obtain *C* cluster representatives 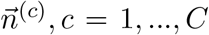, which we call MIDs. To generate robust estimates of the MIDs, we repeat the procedure 100 times on subsampled data sets of size *T/*2 (keeping the ratio of labels) and derive the two quality measures *amount α*^*(c)*^ and *consistency χ*^*(c)*^: As amount we use the fraction of attack vectors 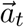 whose normals 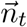 are assigned to the cluster *c* (averaging over all subsamplings). As consistency *χ*^*(c)*^ we denote the fraction of all subsamplings which end up in finding the same cluster *c*. To identify whether MIDs from two susamplings are in the same cluster, we apply the clustering algorithm DBSCAN to all identified MIDs. The performance of DBSCAN can be adjusted by two main parameters. The first parameter is the maximum distance between two vectors in the same cluster, and is set to 0.25 unless mentioned otherwise. The second parameter is the minimum number of samples within a cluster to not be considered as noise, and is set to 3% of the dataset (Ester et al., 1996).

### Bias Correction

MIDs are vectors orthogonal to the decision boundary and thus in order to compute the overlap 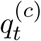 between MID *c* and a specific pattern 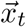, we first subtract the bias 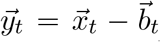 and then compute the scalar products 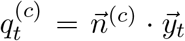. The bias vectors 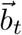 are obtained in every time bin as the center of gravity of the 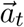 vectors the MID is composed of.

### Relevant Time Bins

To identify whether an activity pattern 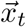 reflects a certain MID, we generate a Null distribution for the overlaps 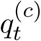 by 1000 random shuffles of the neuron indices. Relevant time bins for MID *c* are those in which 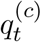 exceeds the upper 97.5%-tile or falls below the lower 2.5%-tile.

### Local field potential analysis

In all recordings from control rats, we selected for the LFP analysis the channels with highest theta power among the tetrodes which were located in the same brain region (CA1 or CA3). Different oscillation bands were extracted by applying a FIR bandpass filter (theta: 6 -11 Hz, slow gamma: 30 - 50 Hz, mid gamma: 55 – 90 Hz, fast gamma: 95 - 140 Hz, ripples: 150 – 250 Hz) based on a Hamming window. Bandpass filtered signals were Hilbert-transformed and the mean square Hilbert-amplitude in a time bin of length *L* was used as an estimate for short-term power analysis.

## Acknowledgements

This work was supported by the German Research Association (DFG) under grant numbers LE2250/13-1 and LE2250/20-1 (FOR 5159) and the NIH under grant number R01 NS086947. The authors also acknowledge support by the state of Baden-Württemberg through bwHPC and DFG through grant no INST 39/963-1 FUGG (bwForCluster NEMO).

## Supplementary Figures

**Figure S1:**
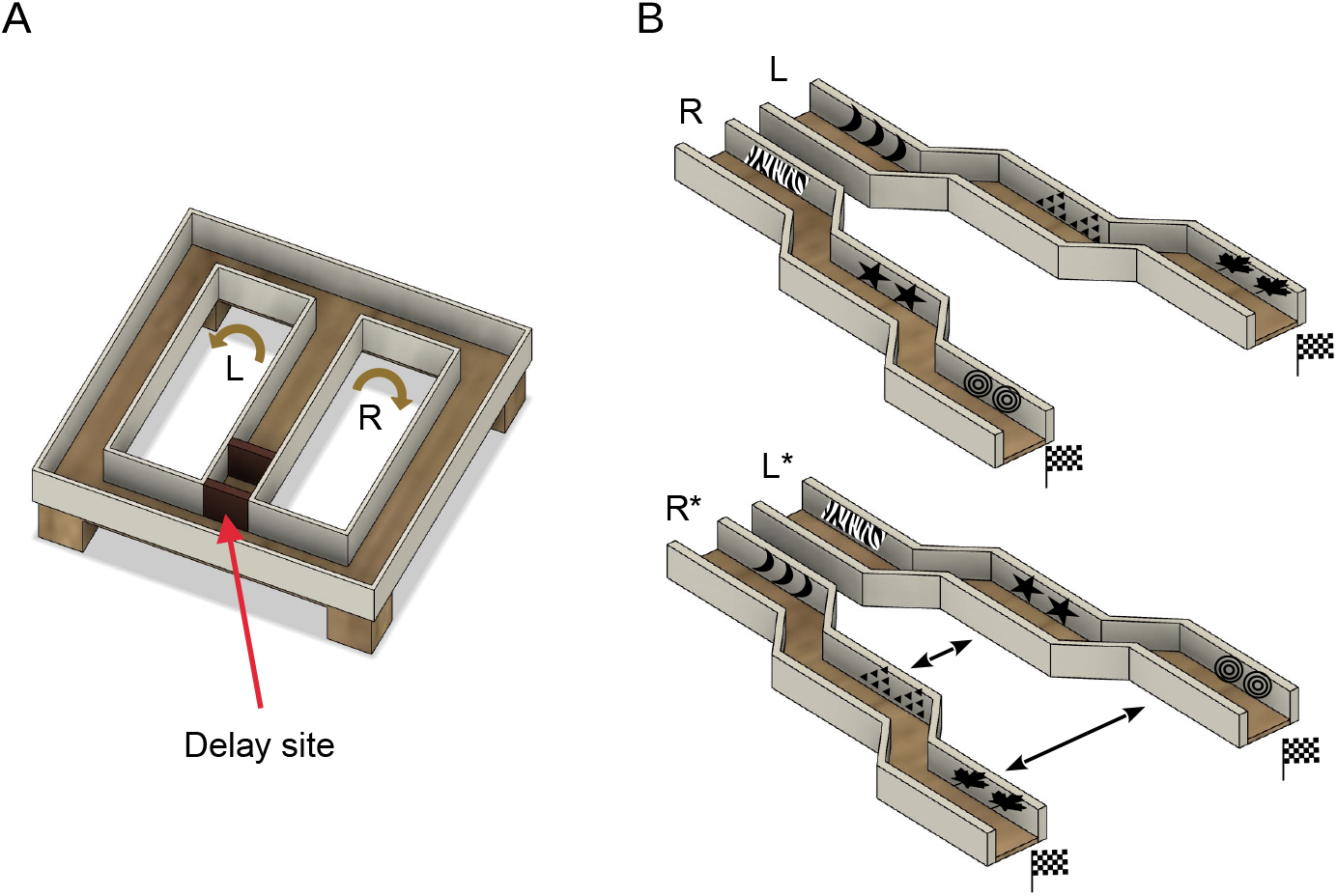
Experimental setup of behavioral tasks. (A) Two groups of rats with and without mEC lesions, are trained to alternate their spatial directional direction in this 8-shaped maze at the end of the middle arm. The rats remain at the delay site for 60 seconds after each trial during which they should maintain a working memory of their previous directional choice. (B) Mongolian gerbils are trained to run on two mazes, in virtual reality, distinguished by left and rightward turns and distinct visual cues placed at the walls of each corridor (top). The gerbils remain stationary for 20 seconds after each maze run during which they receive a reward. Subsequently the gerbils run through previously unseen environments where the visual cues are flipped between the two mazes (bottom).

**Figure S2:**
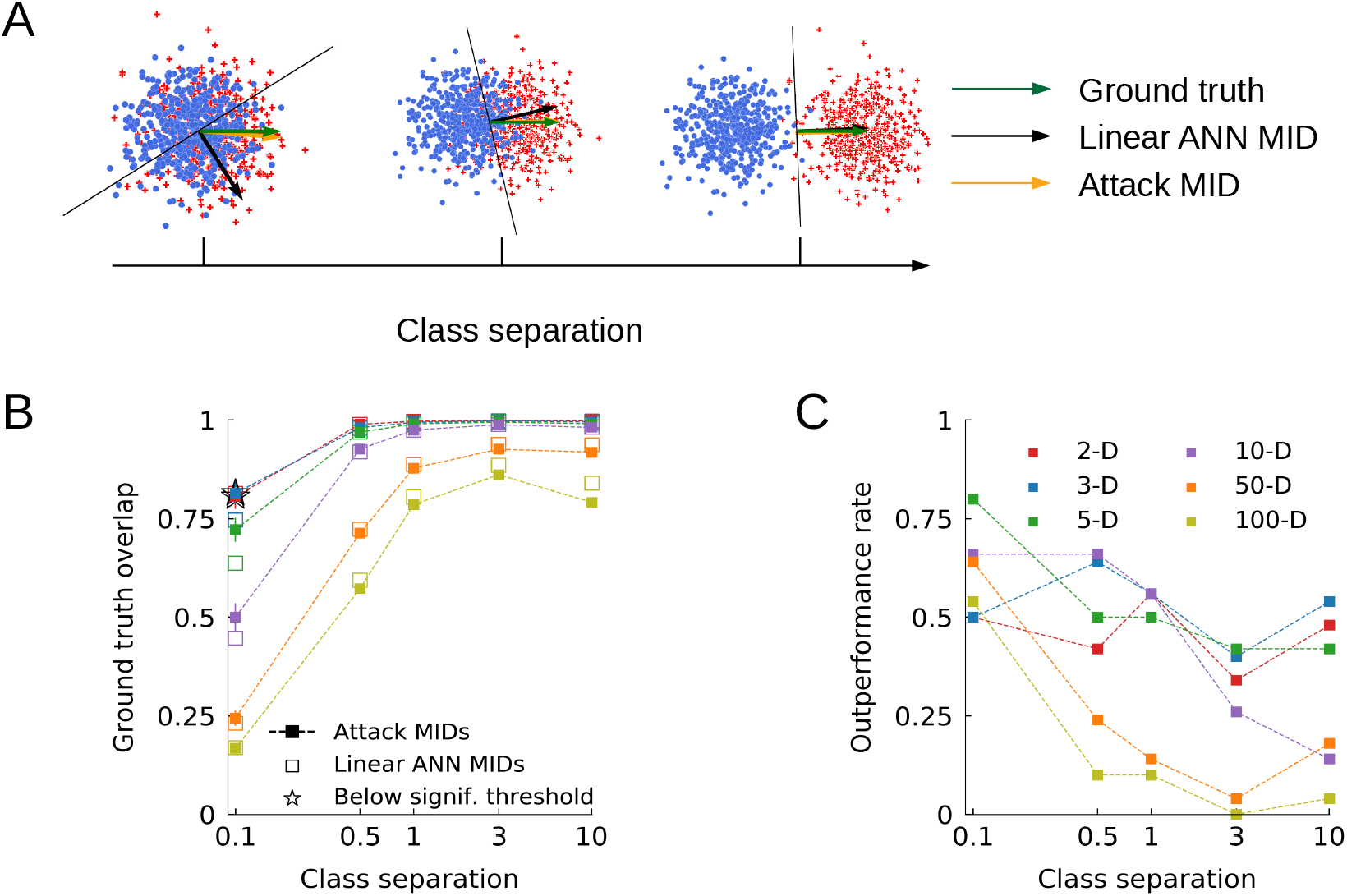
MIDs performance evaluation on artificially generated data sets. (A) Examples of linear binary classification tasks in 2-dimensional space while varying the by-class overlap. MIDs (gold) appear to be closer to the ground truth (green) compared to the ANN weights (black) for high degrees of by-class overlap. (B) Overlap of MIDs and ANN weights with the ground truth for linear classification tasks (*n* = 50). MIDs outperform the linear ANN weight for high dimensionality and high degrees of by-class overlap which is typically the case for neuronal activity. Stars indicate results below significance threshold. The feature space is color coded. (C) Rate at which MIDs outperform the linear ANN weight across repetitions (*n* = 50).

**Figure S3:**
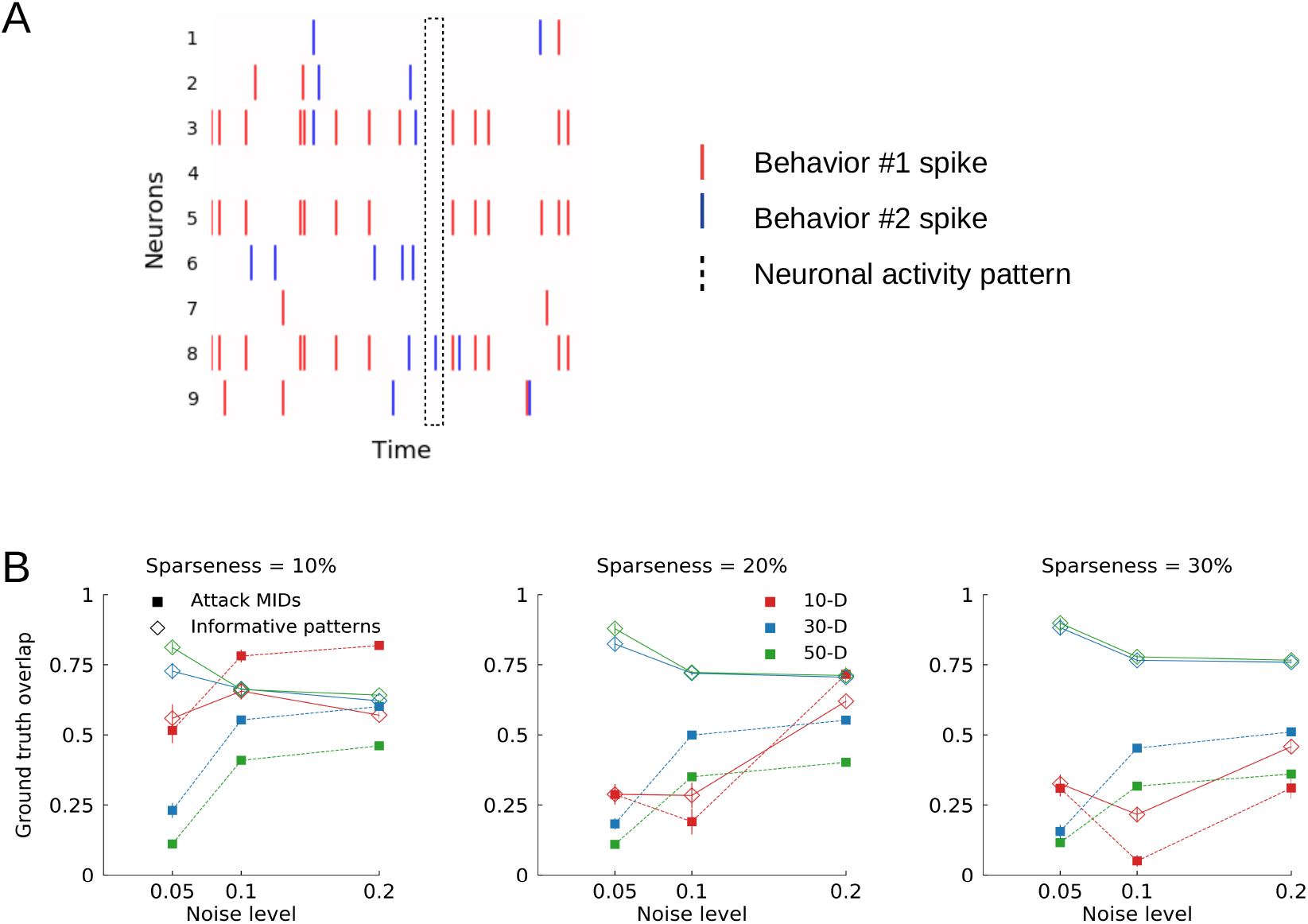
MIDs and informative patterns performance evaluation on artificially generated spiking activity. (A) Schematic of spiking activity for a binary behavioral task. Dotted line indicates a population vector. (B) Overlap of MIDs (square markers) and averaged informative patterns (diamond markers) with the ground truth, while varying the noise, sparseness and amount of noise of the artificial spiking activity. Averaged informative patterns appear to outperform MIDs for activity with similar structure (n=50).

**Figure S4:**
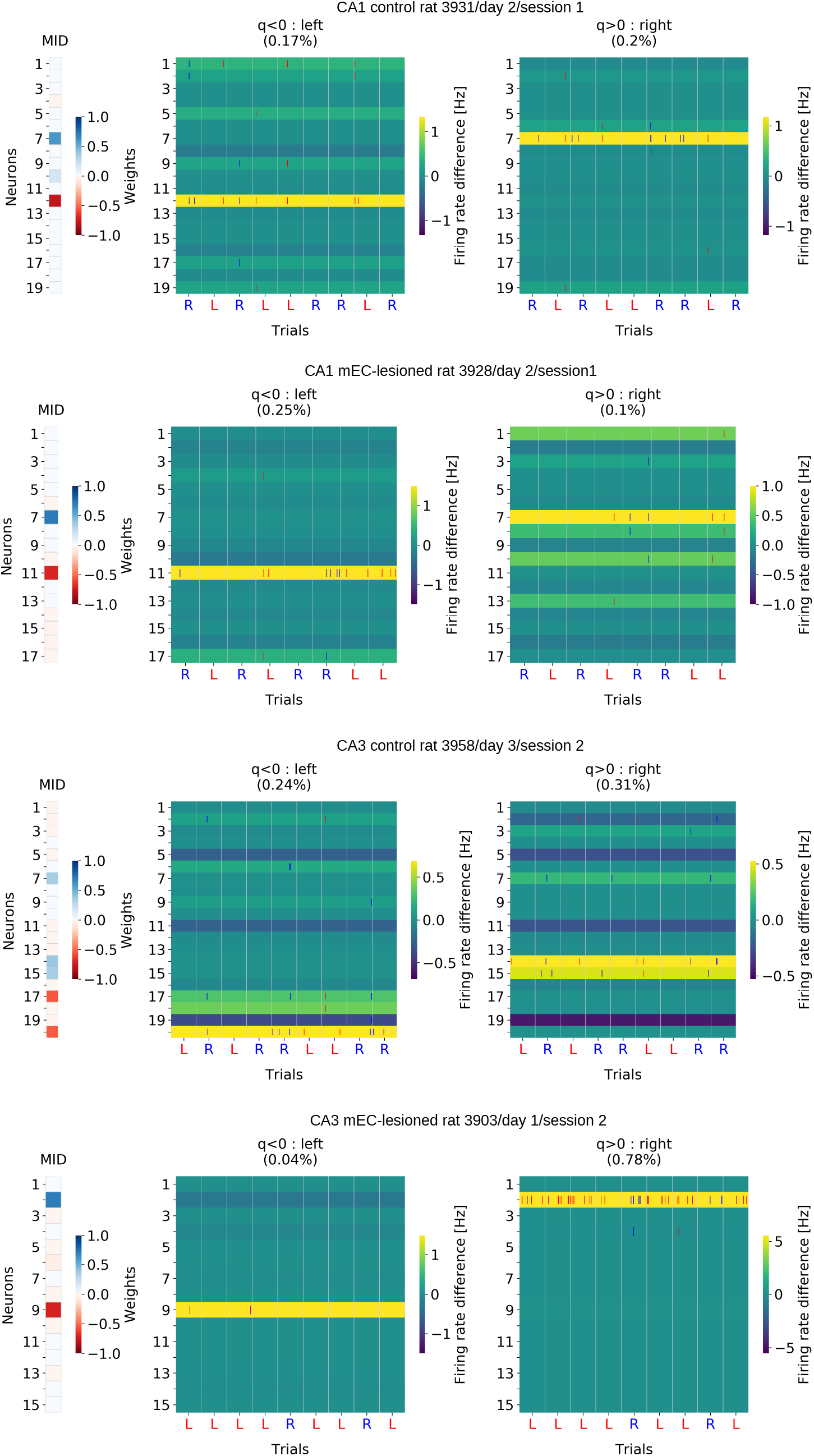
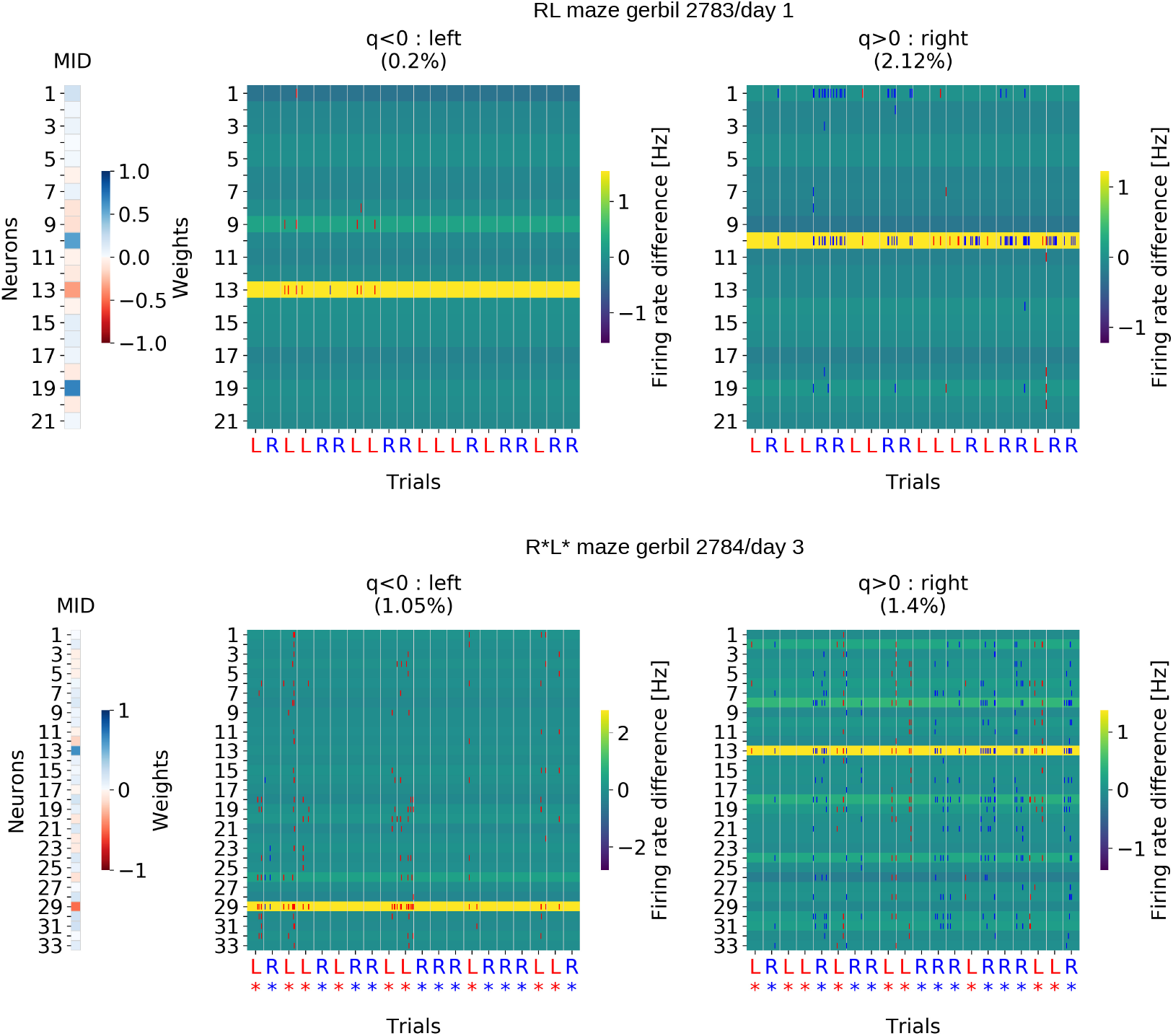
Examples of MIDs (left) and and raster spike plots (right) only for relevant time bins for all groups of experiments. (A) CA1 control rat 3931/day 2/session 1. (B) CA1 mEC-lesioned rat 3928/day 2/session1. (C) CA3 control rat 3958/day 3/session 2. (D) CA3 mEC-lesioned rat 3903/day 1/session 2. (E) RL maze gerbil 2783/day 1. (F) R*L* maze gerbil 2784/day 3.

**Figure S5:**
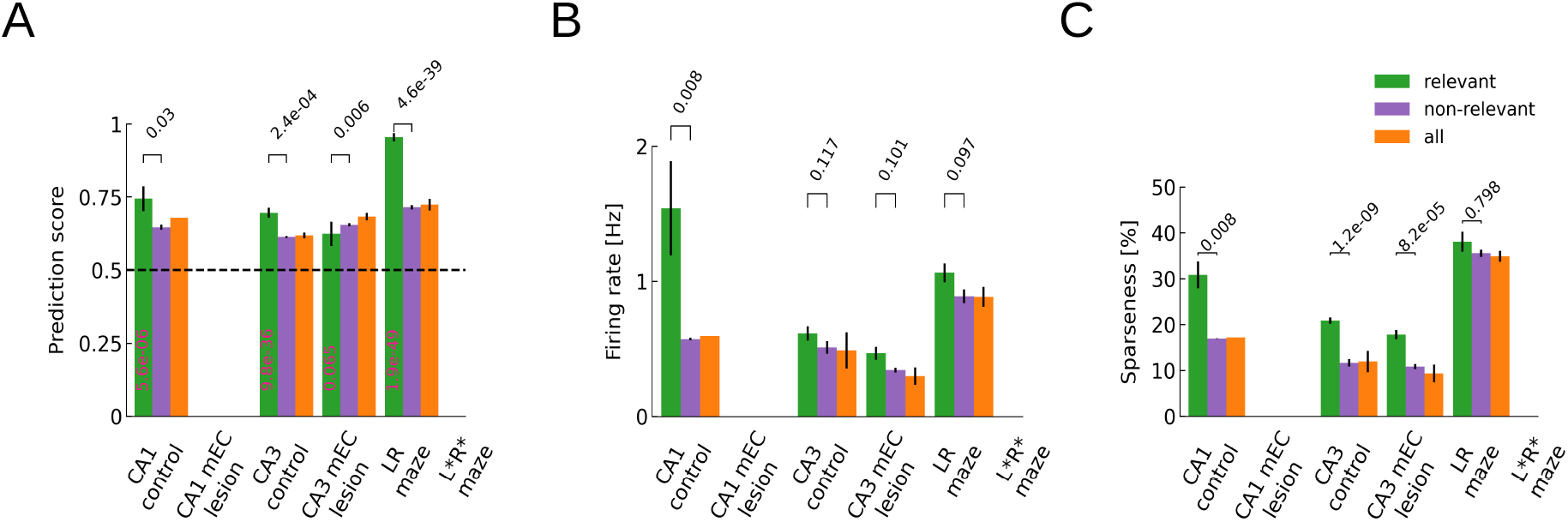
Relevant(green), non-relevant(purple) and all (orange) time bins across significant sessions for *L* = 1000 *ms* time bins. (A) Prediction score comparison. (B) Firing rate comparison. (C) Sparseness comparison.

